# Individualized functional localization of the language and multiple demand network in chronic post-stroke aphasia

**DOI:** 10.1101/2024.01.12.575350

**Authors:** Pieter De Clercq, Alicia Ronnie Gonsalves, Robin Gerrits, Maaike Vandermosten

**Author notes:** **Corresponding author:** Pieter De Clercq. Shared senior authorship (Robin Gerrits, Maaike Vandermosten).

## Abstract

Recent research found a distinct dissociation between brain regions supporting domain-general cognitive processes and regions supporting core language functions. The question of whether individuals with post-stroke aphasia (IWA) exhibit a comparable dissociation remains debated, particularly as previous studies overlooked individual variability in functional network organization and aphasia heterogeneity. To address this gap, we employed an individualized functional localization approach to test the involvement of the domain-general multiple demand (MD) network during language processing in chronic aphasia.

We collected functional MRI data in 15 IWA and 13 age-matched controls. Participants performed a spatial working memory task, triggering MD network activation, as well as a listening and reading task, triggering language network activation. We compared both groups individualized activation patterns and investigated the link with aphasia severity. Involvement of the MD network during language processing was examined by investigating language task activity within subject-specific regions that are active during the MD task.

The language and MD network each generalized well across different task modalities, but exhibited robust spatial dissociation from each other in both groups. Moreover, there was no evidence of MD network activation during language processing in either group. Additionally, the language network showed weaker activation in IWA compared to controls in left-hemispheric brain regions, with higher activation values in the left correlating with improved language performance in IWA.

In conclusion, our findings suggest that the MD network does not contribute to passive, receptive language functions in chronic aphasia or healthy older adults. Instead, our results align with previous research proposing that normalized left-hemispheric language activity supports language performance in chronic aphasia.

## 1 Introduction

Large-scale functional networks within the brain facilitate human behavior, actions and decision-making (Yeo et al., 2011). One prominent network that spans various cognitive processes, such as working memory or executive control, is the domain-general multiple demand (MD) network, involving bilateral frontal and parietal brain regions (Duncan, 2010; Fedorenko et al., 2013). Recent extensive studies in healthy young individuals have demonstrated that the MD network is distinct from the language network (Diachek et al., 2020; Fedorenko and Blank, 2020), which activates a left-lateralized network in temporal and inferior frontal brain areas (Fedorenko et al., 2011). Although the MD network may be involved in language comprehension under specific task demands (e.g., memory probing, semantic judgment), it does not participate in core language functions like passive reading and listening (Diachek et al., 2020) or naturalistic speech listening (Shain et al., 2020).

Populations experiencing language processing difficulties may deviate from typical activation patterns in both the language and MD networks. In individuals with aphasia (IWA), a communication disorder most commonly arising from a stroke in the language-dominant left hemisphere, altered language activation patterns have been observed (Hartwigsen and Saur, 2019; Li et al., 2022; Wilson and Schneck, 2021). For this population, enhanced MD activity (Brownsett et al., 2014; Geranmayeh et al., 2014) and less segregation with the language network (Siegel et al., 2016) have been suggested as a compensation for the neuronal loss.

Prior studies in IWA have monitored the dynamic changes in language reorganization in the brain over time as language abilities recovered. In the acute stage (<1 week) post-stroke, a general reduction in activity in the left-hemispheric language network is observed (Saur et al., 2006; Stockert et al., 2020; Nenert et al., 2018), followed by an upregulation in right hemisphere homotopic language regions during the subacute phase (>1 week to 3 months post-stroke (Saur et al., 2006; Stockert et al., 2020)). In addition to the activation of right hemisphere language regions, MD regions, such as the dorsolateral prefrontal cortex, can become more activated in the subacute phase, with the extent depending on the size and location of the lesion (Stockert et al., 2020). Most research in aphasia focused on the chronic stage (>6 months) post-stroke, characterized by a more stable language profile in IWA (Johnson et al., 2019). Both longitudinal and cross-sectional studies predominantly indicate optimal language recovery through normalization of activation in the left-hemispheric language network (Warren et al., 2009; Crinion et al., 2006; DeMarco et al., 2022; Saur et al., 2006; Stockert et al., 2020; Szaflarski et al., 2011, 2013; Griffis et al., 2017b; Fridriksson, 2010; Pillay et al., 2018; Wilson and Schneck, 2021; Li et al., 2022). The preservation of lesion-free tissue in left hemisphere language regions emerges as a crucial factor in facilitating language recovery (Fridriksson, 2010; Griffis et al., 2017a; Sims et al., 2016; Wilson and Schneck, 2021). Several studies report homotopic right hemisphere recruitment in chronic aphasia as well (Turkeltaub et al., 2011; van Oers et al., 2010; Fridriksson et al., 2009; Griffis et al., 2017b; Postman-Caucheteux et al., 2010; Crinion and Price, 2005; Robson et al., 2014), but it has been associated with both better (van Oers et al., 2010; Griffis et al., 2017b; Robson et al., 2014) and worse (Postman-Caucheteux et al., 2010; Szaflarski et al., 2013) behavioral outcomes (for a recent overview, see Li et al. (2022)). Similarly, multiple studies found involvement of what they considered MD brain regions when IWA performed language tasks (Allendorfer et al., 2012; Brownsett et al., 2014; Geranmayeh et al., 2014), but this has been suggested as both an adaptive (Geranmayeh et al., 2017) and maladaptive (LaCroix et al., 2021) mechanism to support language recovery.

A systematic review and meta-analysis on neuroplasticity in aphasia concluded that there is in principal only minimal evidence supporting the recruitment of MD regions or other brain regions outside the language network (Wilson and Schneck, 2021). However, the review noted that previous studies lacked methodological robust comparisons against controls, and that the reported MD regions associated with language performance in IWA were often implicated in healthy controls as well for similar task demands. Consequently, the nature of observed activation in MD regions as a true compensatory mechanism and its implication in language recovery are questionable. Moreover, prior studies in aphasia were restricted to traditional group-based anatomical region of interest (ROI) analyses, relying on the critical assumption that a given anatomical brain region corresponds to the same functional units for all individuals. Yet, research has shown that this assumption is fallacious (Blank et al., 2017) and neglects inter-individual variability in both the function and anatomy of brain regions (Juch et al., 2005; Fedorenko et al., 2010, 2012; Blank et al., 2017). As a consequence, there is a substantial reduction in sensitivity and functional resolution in the analyses (Fedorenko et al., 2010; Shain et al., 2020), which could be even more detrimental for aphasia research (Blank et al., 2017), given the large heterogeneity in lesion characteristics and individual recovery patterns (Jarso et al., 2013; Sebastian et al., 2016; Li et al., 2022). For these reasons, prior researchers have advocated to use the variability across participants as a variable of interest rather than measurement noise (Blank et al., 2017; Seghier and Price, 2018).

As an alternative to group-based averaging, functional localization adopts an individualized approach for selecting ROIs (see Fedorenko et al. (2010)). This method selects functional ROIs based on individual activation patterns in localizer tasks performed by the participant. Language localizer tasks, such as passive visual reading (Fedorenko et al., 2010) or auditory listening (Scott et al., 2017), have been implemented in big-sample studies and validated across multiple languages (Malik-Moraleda et al., 2022). The MD network generalizes well across many different localizer tasks engaging cognitive effort, e.g., spatial working memory, inhibitory control or arithmetic computations (Fedorenko et al., 2013). Although both networks can often lie in close proximity to each other, particularly in frontal brain regions like the inferior frontal gyrus and Broca’s area (Fedorenko et al., 2012), each can activate distinct subregions of the same anatomical brain area and exhibit robust functional and spatial dissociation (Diachek et al., 2020). Whether older individuals or IWA experiencing language processing difficulties exhibit such dissociation, remains to be tested with functional localization. Yet, this technique has the potential to investigate the involvement of the MD network during language processing in a more sensitive, statistically powerful and individualized approach.

In the present study, we applied subject-specific functional localization to examine the language and the MD network in chronic post-stroke aphasia and healthy older individuals. To identify the language network, we administered both a passive auditory listening (Scott et al., 2017) and a passive visual reading task (Fedorenko et al., 2010). For MD network localization, a visual spatial working memory task, known to engage the MD network in healthy young adults (Fedorenko et al., 2013), was utilized. Our first objective involved comparing the activation magnitude and spatial distribution of the networks in both groups. Secondly, we aimed to explore the involvement of subject-specific MD networks during language processing, with a particular focus on potential increased activation in IWA. For both objectives, we explored differential effects of the two hemispheres, investigating the role of the right hemisphere as a potential compensation mechanism in aphasia. Finally, we assessed the robustness of individually-selected networks by cross-examining the activation of selected voxels across localizer tasks.

## 2 Materials and methods

### 2.1 Participants

Our sample comprised 15 IWA (5 female, 65 *±*16 y/o) in the chronic phase post-stroke (*≥* 12 months) and 13 neurologically healthy controls (4 female, 69 *±*7 y/o). No significant age difference was present between groups (unpaired Wicoxon rank sum test: W=89.5 , p=0.73). 12 IWA were recruited at the stroke unit of the University Hospital Leuven in the acute stage post-stroke, and were invited to participate in the fMRI study in the chronic stage. The other 3 IWA were recruited via speech-language pathologists. Healthy controls were recruited making sure they matched the age of IWA at the group level, and making sure they matched the number of left-handers (2 in both groups, as assessed with the Edinburgh Handedness Inventory, Oldfield (1971)). The inclusion criteria for IWA were: (1) a left-hemispheric or bilateral stroke, (2) a diagnosis of aphasia in the acute stage after stroke using behavioral language tests and (3) no formal diagnosis of a psychiatric or neurodegenerative disorder. For more information regarding recruitment strategy and diagnosis in the acute stage post-stroke, we refer to (Kries et al., 2023), where 12 of our 15 IWA also participated. Furthermore, a detailed overview including demographic information (age, sex, handedness, time since stroke, speech-language therapy, performance on diagnostic tests for aphasia at time of participation) and lesion information (stroke type, affected blood vessel, lesion size and lesioned hemisphere) about the aphasia sample can be found in the Supplementary Materials Table 1. For both groups, only participants without diagnosed hearing disabilities were included as they needed to listen to speech stimuli in the scanner. The study was approved by the ethical committee UZ/KU Leuven (S60007), and all participants gave written consent before participation. Research was conducted in accordance with the principles embodied in the Declaration of Helsinki and in accordance with local statutory requirements.

**Table 1.** Distribution of the selected voxels across the predefined parcels.

|  | **Listening** |  | **Reading** |  | **spWM** |  |
| --- | --- | --- | --- | --- | --- | --- |
|  | Controls | IWA | Controls | IWA | Controls | IWA |
| Language parcels | 61.43% | 66.87% | 66.46% | 62.77% | 7.10% | 11.02% |
| MD parcels | 29.91% | 26.03% | 28.96% | 31.78% | 89.12% | 85.19% |
| Member of both (MFG, IFG) | 8.66% | 7.10% | 4.58% | 5.45% | 3.78% | 3.79% |

All participants completed standardized clinical tests for aphasia at the time of participation, namely the ’Nederlandse Benoemtest’, i.e., Dutch Naming Test (Van Ewijk et al., 2020), and the ScreeLing Test, i.e., a comprehensive aphasia battery testing for phonological, semantic and syntactic language skills (Visch-Brink et al., 2010; El Hachioui et al., 2017). IWA scored significantly lower on both behavioral tests compared to the control group (Dutch Naming Test: W=15, p<.001; ScreeLing: W=23.5, p<.001). Furthermore, the Oxford Cognitive Screen-NL was administered to assess cognitive functioning (Huygelier et al., 2019). We used a composite score of the subscales attention, executive functioning and memory to measure cognitive functioning. For more information on these tests, we refer to (Kries et al., 2023). We found no group difference on this test (W=86, p=0.61).

### 2.2 MRI data acquisition

Participants completed an MRI protocol consisting of a T1-weighted anatomical scan, a T2-weighted FLAIR scan, and 4 functional paradigms: an auditory listening localizer task, a visual reading localizer task, a spatial working memory (spWM) localizer task, and a natural story listening paradigm. The latter paradigm will not be considered in the present study. In total, the entire scan protocol lasted 50-60 minutes.

All MRI data were collected on a 3-T Philips scanner with a 32-channel head coil at the University Hospital Leuven. A high-resolution anatomical image of the brain was acquired using a T1-weighted MPRAGE sequence (TR9.687ms, TE=4.6ms, 182 sagittal slices, flip angle=8ř, isotropic voxel size=1mm). The T2-weighted FLAIR scan (TR4800ms, 200 sagittal slices) was also obtained and used to identify and segment lesions. Functional images were acquired using a T2*-weighted single-shot EPI sequence (TR1300ms, TE=29.8ms, 52 near-axial slices, flip angle=90ř, voxel size=187x1.87x2.7mm). During functional imaging, stimuli were presented via a translucent screen at the back of the magnet bore, which participants saw via a mirror attached to the head coil.

### 2.3 Lesion segmentation

Lesion masks were manually segmented on each patient’s T2-weighted FLAIR images using MRIcron, guided by the report in the patient’s medical file (written by a neurologist/neuroradiologist). A lesion overlap map of our aphasia sample is provided in Figure 1. Largest lesion overlap was found in the left insula and left inferior frontal gyrus (overlap of 10/15 participants).

**Figure 1.**
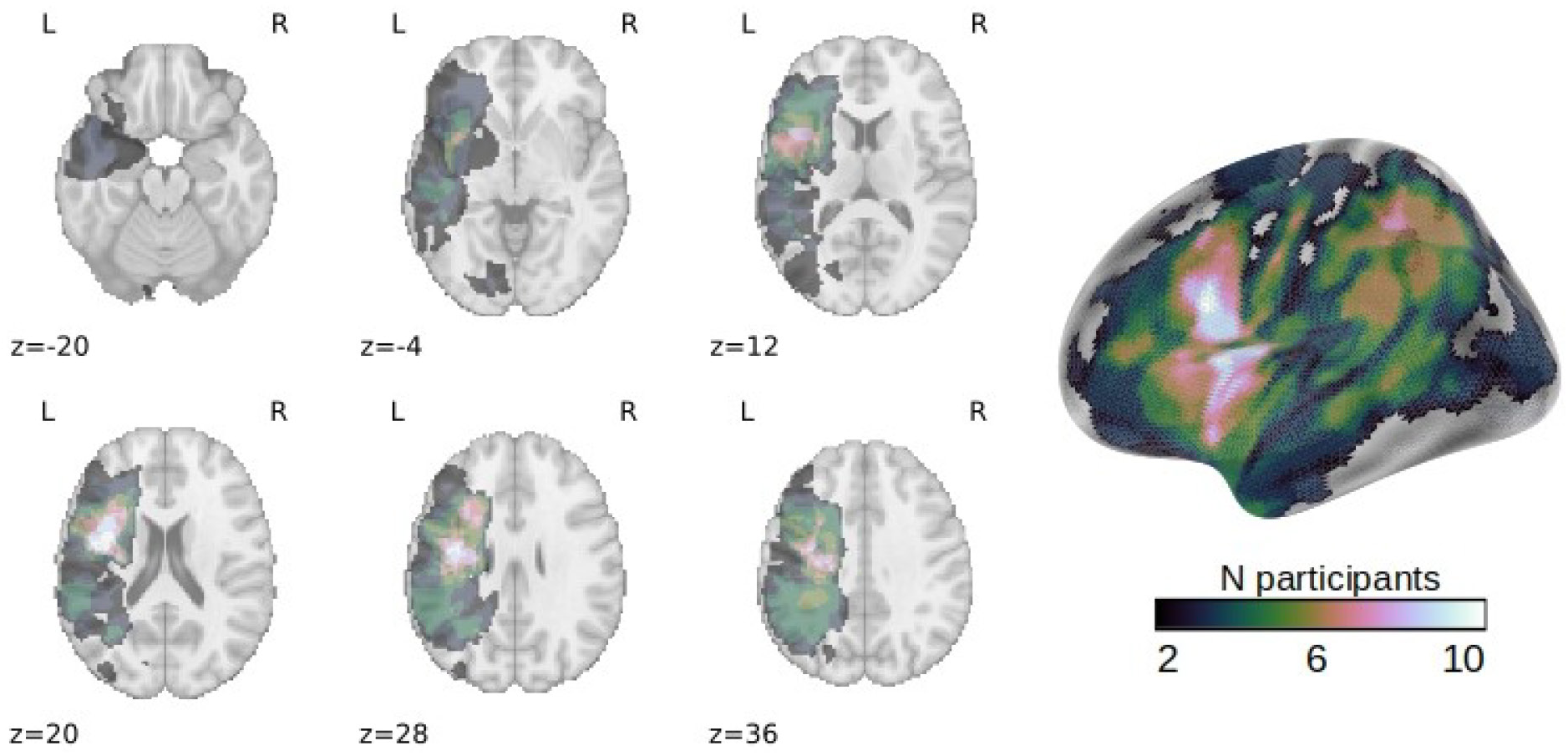
Lesion overlay image. Axial slices with corresponding Z-coordinates are shown in neurological orientation. On the right, a surface image is shown.

### 2.4 fMRI tasks

We aimed to localize the language network using both an auditory listening and a visual reading task, and the MD network using a visual spatial working memory task (spWM). These tasks are described below.

#### Listening localizer task

The listening localizer was implemented following the methodology outlined by (Scott et al., 2017), demonstrating robust language network localization in healthy, young listeners. We extracted 20 segments of 18 seconds each of the fairytale “De Kleine Zeemeermin” (“The Little Mermaid”, written by Christian Andersen and narrated in Dutch by a male Flemish-native speaker), which we used in prior work in our research lab (Accou et al., 2021). We created acoustically degraded versions of 10 segments, which were used as a control condition to the other remaining 10 segments, devoid of any noise or filtering. The degraded versions were generated by introducing noise with the same spectrum as the intact stimuli, ensuring that the degraded condition retained identical acoustical and spectral information while lacking speech understanding and linguistic processing. The modulator for these stimuli was derived from the envelope of the intact speech, calculated as the absolute value of the Hilbert transform. We also included a gradual volume fade-out of 2 seconds at the end of each segment and 8 rest blocks of 12s where no auditory stimuli were presented, consistent with Scott et al. (2017); Malik-Moraleda et al. (2022). Condition order was randomized, but we made sure that two identical conditions never appeared consecutively. In total, the listening localizer lasted 7 minutes and 36 seconds. Prior to entering the scanner, participants performed practice trials to familiarize with the task, and audibility of the stimuli was ensured through an fMRI dummy scan. Participants were instructed to listen attentively to all stimuli while maintaining fixation on a centrally presented cross on the screen. All participants performed this task.

The contrast in activation “intact>degraded” served as the primary focus to localize the language network. This contrast exhibits stronger responses in language-specific brain regions, as shown in previous research (Scott et al., 2017; Malik-Moraleda et al., 2022). These regions are specifically involved in processing word meanings and combinatorial semantic/syntactic processes, which are not activated when processing the degraded stimuli.

#### Reading localizer task

The reading localizer included two critical conditions: passively reading sentences and reading a list of non-words. We created the stimuli consistent with prior research (Fedorenko et al., 2010), but we adapted task difficulty (i.e., slower presentation and shorter sentences) to match our target population (i.e., elderly and IWA). The sentences condition consisted of simple syntax structures, where word-level meanings are combined into larger phrase-level and sentence meanings (e.g., “ALL EMPLOYEES RECEIVED A NICE BONUS LAST YEAR”). The non-words consisted of pronounceable pseudowords that matched the length of the words in the sentence condition. non-words were generated with the Wuggy pseudoword generator (Keuleers and Brysbaert, 2010). Each trial consisted of a sentence or word list of 8 (non)words, each presented for 1000ms, and a cue that the sentence had ended for 1000ms (a green check mark). In total, each trial was 9 seconds long, and a condition block consisted of 2 consecutive trials of the same condition (i.e., 18 seconds in total). The reading localizer task included 10 blocks for both conditions and 8 rest blocks of 12 seconds each (where participants had to fix their eyes on a fixation cross), with randomized order (with the constraint that two identical conditions never appeared consecutively). In total, the reading localizer task lasted 7 minutes and 36 seconds. Participants were instructed to silently read the sentences and non-words. Participants also underwent task familiarization before going into the scanner (i.e., silently reading practice trials), and readability of the sentences was ensured through an fMRI dummy scan. The procedure of this task is visualized in the Supplementary Materials, Figure 1. We had missing data for 2 healthy controls and 1 IWA because of limited scanning time (two participants) or having to stop the scan early due to back pain from lying in the scanner (one participant). In total, 14 IWA and 11 heatlhy controls successfully performed the reading localizer task.

The contrast in activation “sentences>non-words” was used as the critical contrast to localize the language network. Language-specific brain regions are known to exhibit stronger responses to meaningful sentences (Fedorenko et al., 2010), as they are engaged in semantic and syntactic processes, which are not activated when reading non-words.

In addition, the contrast of “non-words>sentences” served a secondary focus. Given that processing non-words is more cognitively demanding, the contrast “non-words>sentences” is frequently employed in the literature on healthy young participants to localize the MD network (e.g., see (Fedorenko et al., 2013; Shain et al., 2020; Diachek et al., 2020)). We did not use this contrast to localize the MD network in our main analysis (i.e., Analysis 1 and 2), because this approach would result in a spatial dissociation between the MD and language networks as defined by the reading localizer a priori (i.e., activation in a certain voxel would indicate higher activation for either one of these two contrasts). However, we did use this contrast to test the reliability and generalizability of the defined MD network in Analysis 3.

#### Spatial working memory localizer task

We administered a visual spWM task contrasting a harder condition with an easier condition, following the methodology established in prior research (Fedorenko et al., 2013). Yet, the procedure was adapted to suit our target population, with a slower pace and a reduced number of presentations. On each trial (9 seconds), participants saw a fixation cross for 500 ms, followed by a 3x4 grid where either one (easy condition) or two (hard conditions) locations were flashed for 2250 ms, twice. Subsequently, participants mentally combined the flashed locations and responded to a two-alternative (left/right) forced-choice paradigm, shown during a 4000ms window. Participants held buttons in their left and right hands for responses. In cases of paralysis on one side (N=2 IWA), individuals with paralysis used a single button in one hand, incorporating left and right options on the same button. Each condition block comprised 2 consecutive trials of the same condition (totaling 18 seconds). The task comprised 8 blocks per condition, interspersed with 6 rest blocks lasting 12 seconds each, during which participants fixed their eyes on a central cross. Condition order was randomized, with a constraint to prevent consecutive appearances of identical conditions. In total, the task duration was 5 minutes and 48 seconds. Participants underwent a training phase outside the scanner, and visibility of the stimuli was ensured through an fMRI dummy scan. The procedure of the task and behavioral outcomes are provided in the Supplementary Materials, Figure 1. We had missing data for 1 IWA due to limited scanning time, totalling 14 IWA and 13 heatlhy controls who successfully performed the spWM localizer task.

The “hard>easy” contrast served as the critical contrast, which was previously found to robustly activate the MD network (Fedorenko et al., 2013). This network is associated with cognitively and atten- tively demanding tasks and exhibits generalization across various other demanding cognitive activities (e.g., arithmetic operations, see Fedorenko et al. (2013)).

### 2.5 fMRI data analysis

fMRI data analysis was conducted using SPM12 (Welcome Trust Centre for Neuroimaging, London, UK), implemented in Matlab 2021b (MathWorks, Massachusetts, USA). Preprocessing included motion correction through rigid realignment, followed by slice-timing correction. Subsequently, the functional images were coregistered to the participant’s T1 image, utilizing the mean functional image. Normalization was then performed to the template for older individuals in MNI space with the Clinical Toolbox (Rorden et al., 2012), taking into account the lesion and intact contralateral hemispheric brain regions in normalizing the T1 image. Finally, functional images underwent smoothing using a Gaussian kernel with a size of 4 mm.

Next, first-level analyses were conducted with each participants preprocessed volumes. Experimental conditions were modeled by convolving with the canonical hemodynamic response function (Friston et al., 1994), and beta weights were estimated using a generalized linear model. Six estimated head motion parameters (i.e., translation and rotation in the x, y, and z directions) were included as nuisance variables in the model. Contrasts of interest for each task were obtained by comparing task-related activity against activation in the control condition: “intact>degraded” (listening), “sentences>non-word”s (reading), and “hard>easy” (spWM).

#### Subject-specific network selection

We applied subject-specific functional localization to identify the language and MD network. This technique selects regions of interest (ROIs) based on the participants activation patterns in the localizer task. By contrast, traditional fMRI experiments in aphasia research typically select ROIs anatomically, which is illustrated in Figure 2A. However, this approach does not account for inter-subject variability in functional network organization, resulting in reduced sensitivity and functional resolution (Fedorenko et al., 2010). An example of inter-subject variability in the language network for 2 IWA and 2 healthy control participants in our sample is provided Figure 2B.

**Figure 2.**
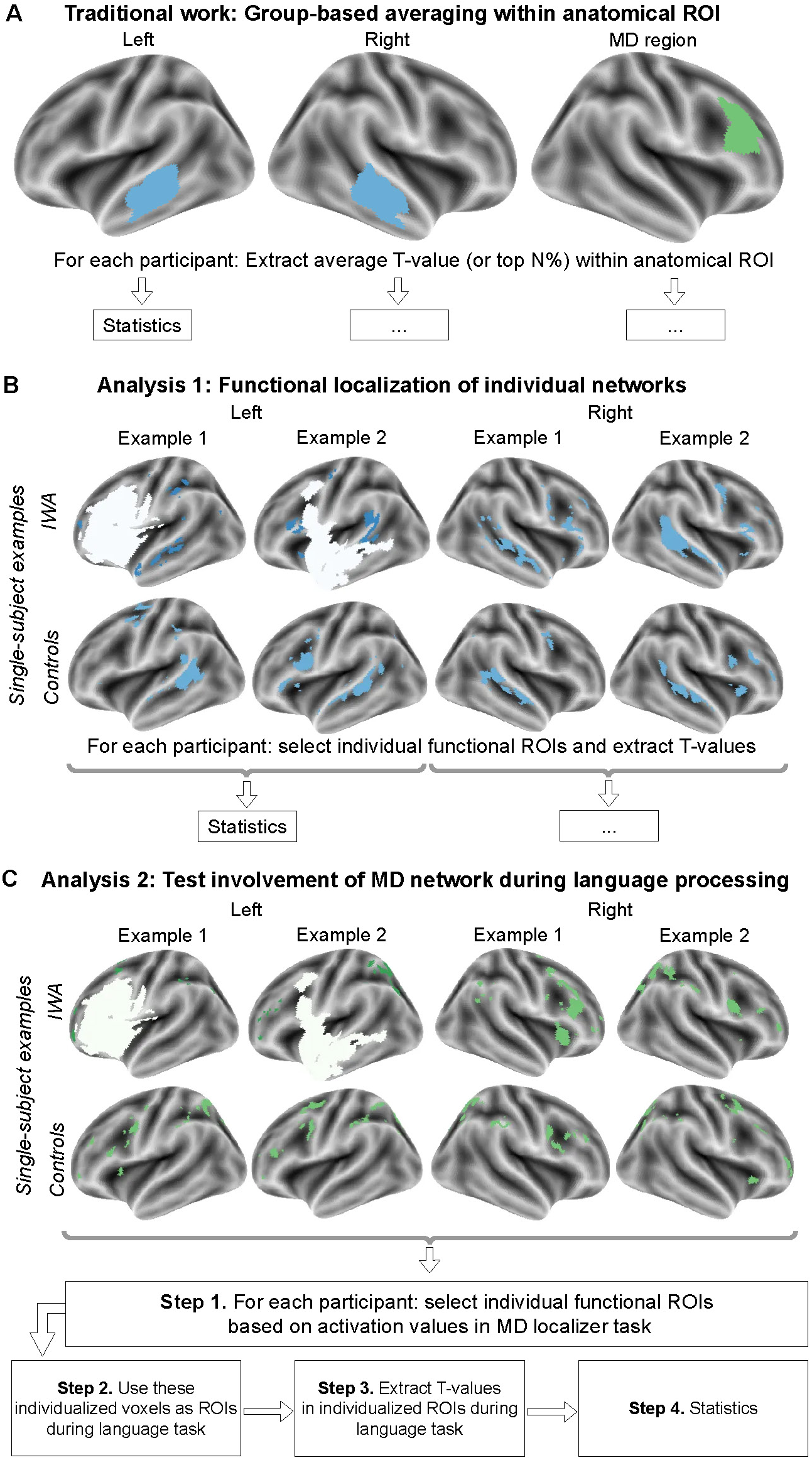
Methodological overview of functional network selection and statistical analyses. **A.** Traditional fMRI research in aphasia applies group-based averaging in pre-defined anatomical ROIs. Examples of single-subject language network selection for Analysis 1, defined as the top 10% most active voxels across the union of language and MD parcels for the contrast of interest (applying T-value and cluster-thresholding). This example, showcasing variability in lesion (white, IWA) and functional organization of the language network, involves the listening localizer task, but the same approach was applied for the reading localizer task and spWM localizer task for MD network selection. **C.** Visualization of our approach to investigate the involvement of the MD network in Analysis 2: Activation during language tasks is investigated in individually-defined voxels during the spWM localizer task. In the Supplementary Materials, results within-isolated parcels (not applying thresholding) are presented, similar to prior work (Diachek et al., 2020; Shain et al., 2020).

The language network was localized using critical language contrasts in both the listening and reading localizer tasks. While both localizers are believed to reveal the same modality-independent language network (Scott et al., 2017), this assertion is based on studies involving healthy young individuals and has not been tested in healthy elderly and stroke patients. Therefore, one of our aims was to assess the generalizability of results across modalities and examine the similarity between the language networks derived from the listening and reading localizers. The MD network was localized using the contrast in activation between the hard and easy conditions in the spWM localizer task.

The selection of subject-specific networks was based on methodologies established in prior research work (Diachek et al., 2020; Shain et al., 2020). The procedure involves two main steps: 1) applying group-level masks to the individual T-maps (parcels; available for download from https://evlab.mit.edu/funcloc/ download-parcels) and 2) selecting the top 10% most active voxels within the parcels. The language-specific parcels are based on a large group study involving 220 young participants and include 6 bilateral parcels (i.e., 12 in total): the inferior frontal gyrus (IFG) and its orbital part (IFGorb), middle frontal gyrus (MFG), anterior temporal cortex (AntTemp), posterior temporal cortex (PostTemp), and angular gyrus (AngG). The MD-specific parcels are based on a large group study involving 197 participants who completed the spWM localizer task and include 10 bilateral masks (i.e., 20 in total): in the posterior (PostPar), middle (MidPar), and anterior (AntPar) parietal cortex, precentral gyrus (PrecG), superior frontal gyrus (SFG), middle frontal gyrus (MFG) and its orbital part (MFGorb), opercular part of the inferior frontal gyrus (IFGop), the medial prefrontal cortex (mPFC) and the insula (Insula). Overlap between language and MD parcels are present in parts of the MFG and IFG. The parcels are displayed on a surface map in the Supplementary Materials, Figure 2.

In prior research, the (unthresholded) top 10% most active voxels within all isolated parcels were selected (e.g., Shain et al. (2020); Diachek et al. (2020), and voxel selection was confined to language parcels for the language network and vice versa for the MD network. However, this approach ensures spatial dissociation between the language and MD networks a priori, given their involvement in spatially dispersed ROI parcels. Therefore, we modified this procedure and opted to select the top 10% most active voxels across the union of all (language + MD) parcels. To prevent the inclusion of singular, noisy voxels in a participants network, we applied T-value and cluster thresholding using a T-value threshold corresponding to p<0.05 (*T ≈* 1.65) and a cluster threshold of 20. Some exemplary results of voxel selection are presented in Figure 2B and C. Additionally, we also present results within-isolated parcels as typically performed in prior research (top 10%, unthresholded) in the Supplementary Materials to offer a comprehensive overview compared to traditional functional localization approaches.

### 2.6 Statistical Analysis

#### Analysis 1: spatial distribution and activation of networks

##### Spatial distribution

First, we characterized the spatial distribution of the networks, defined as the top 10% most active voxels across the union of all parcels (including both language and MD parcels, visualized for example participants in Figure 2). We provided network overlap maps for all three localizer tasks, and detailed the spatial distribution of the selected voxels across the language and MD parcels. Based on prior literature, we expected most selected voxels to be located within the language parcels for the language tasks and within the MD parcels for the MD task (spWM).

Subsequently, we investigated the spatial overlap (i.e., the similarity) between the different networks as measured by the Dice score:

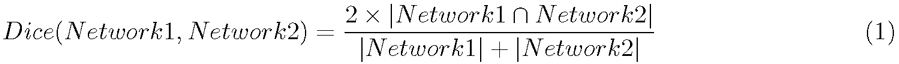

The Dice score provides a value between 0 (indicating no spatial overlap, i.e., no shared voxels) and 1 (representing complete spatial overlap). This calculation was performed within each individual subject for three comparisons: 1) between the listening and reading network, 2) between the listening network and the MD network, and 3) between the reading and MD network. With this analysis, our aim was to quantify both 1) the extent to which the reading and listening networks share similar spatial regions, and 2) whether the brain utilizes the same spatial regions for both language processing and performing cognitively demanding tasks (i.e., the spWM task). Group differences (controls vs. IWA) were explored using non-parametric Wilcoxon rank-sum tests.

##### Network activation in anatomical parcels

We also conducted comparisons in activation magnitude (i.e., T-values) between parcels and between participant groups for all 3 localizer tasks. Specifically, we identified the top 10% most active voxels separately across the language parcels and MD parcels for each hemisphere (i.e., 4 parcels, namely left and right language, and left and right MD). We splitted the analyses for both hemispheres, as the role of the right hemisphere in aphasia remains a subject of debate (Price and Crinion, 2005; Heiss and Thiel, 2006; Crosson et al., 2007; Li et al., 2022; Wilson and Schneck, 2021). However, we noticed that for some participants and some parcels (i.e., in some of these 4 parcels), no voxels survived thresholding. Therefore, T-value thresholding was decreased to 0 for the purpose of this analysis.

First, we investigated whether language parcels exhibited higher T-values during language tasks compared to MD parcels (irrespective of hemisphere and group), and vice versa. To this end, we ran a repeated measures ANOVA analysis per localizer task with the following linear mixed effects model:

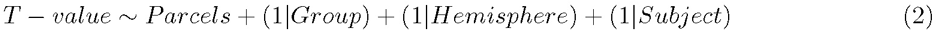

Second, we explored group differences in all 4 parcels (union in language and MD, left and right) using Wilcoxon rank-sum tests. Exploratory analyses within isolated parcels (unthresholded top 10% active voxels per isolated parcel) are provided in the Supplementary Materials.

#### Analysis 2: subject-specific MD network activation during the language localizer tasks

Next, we investigated the involvement of the subject-specific MD network during language processing. While the Dice score between the language and MD network in Analysis 1 may indicate the extent to which both networks share brain regions, it does not directly assess whether the subject-specific MD network is actually activated during language processing. To address this, we first selected the subject- specific MD network revealed by the spWM localizer task (i.e., the top 10% most active voxels across MD parcels using thresholding). Subsequently, we used this subject-specific MD network as ROIs during the listening and reading localizer tasks to test whether the activation in these ROIs was significantly larger than 0 (one-sided Wilcoxon signed-rank tests). In the event of significant activation in these ROIs, group comparisons were conducted using Wilcoxon rank-sum tests (p-values FDR-corrected). This approach of single-subject MD network investigation during language processing is identical to prior research in healthy young participants (see (Diachek et al., 2020)) and is illustrated in Figure 2C. Exploratory analyses within isolated parcels (unthresholded top 10% active voxels per isolated parcel) are provided in the Supplementary Materials.

#### Analysis 3: Reliability of selected networks using across-task cross-validation

Finally, we assessed whether individually-selected networks are reliably defined and whether they gener- alize over task modalities using an across-task cross-validation approach. To do so, a similar approach as Analysis 2 was applied: we first selected the subject-specific language network of a first language contrast (top 10% voxels across language parcels in the e.g., “intact>degraded” contrast during the listening task) and tested its activation for the second language contrast (“sentences>non-words” during the reading localizer), and vice-versa. Similarly, we tested the generalizability of the MD network by first selecting the subject-specific MD network of a first MD contrast (top 10% voxels across MD parcels in the e.g., “hard>easy” contrast during the spWM task) and tested its activation for the second MD contrast (“non- words>sentences”), and vice-versa. Testing activation was identical to Analysis 2, namely testing whether observed T-values at group level were significantly higher than 0 using one-sided Wilcoxon signed-rank tests. As outlined above, the “non-words>sentences” contrast was not used in the prior 2 Analyses as it would result in a spatial dissociation between the language and MD network as defined in the reading localizer a priori. Nevertheless, this contrast is frequently used in the literature in healthy young partic- ipants to localize the MD network as well (e.g., (Diachek et al., 2020; Shain et al., 2020)), and here we test whether the MD network as defined using the spWM task also generalizes to the reversed language contrast in older individuals and IWA.

Our across-task cross-validation approach is similar to an across-runs cross-validation approach as frequently employed in studies testing healthy young participants (e.g., (Diachek et al., 2020; Shain et al., 2020)). Yet, instead of cross-validation across runs of the same task, we apply cross-validation across different tasks, which addresses one of our research objectives described in the introduction to test whether our results generalize across different tasks and task modalities, as suggested in prior work involving healthy young participants (Fedorenko et al., 2013; Scott et al., 2017; Diachek et al., 2020).

##### Link to behavioral performance

We explored the link between activation patterns and aphasia severity (ScreeLing test score) within the aphasia group. Specifically, we computed partial Spearman rank correlations between network activation and the ScreeLing score, correcting for age, lesion size and the time since stroke. P-values were FDR- corrected for multiple comparisons.

## 3 Results

### 3.1 Analysis 1: Spatial distribution and activation of networks

#### Spatial distribution

We begin by detailing the spatial distribution of networks, defined as the top 10% most active voxels during language localizers (listening and reading) and the MD localizer (spWM task) across the union of all parcels. An overlap map, visualizing the overlap across subjects per localizer task with an overlap threshold of 20%, is provided in Figure 3A.

**Figure 3.**
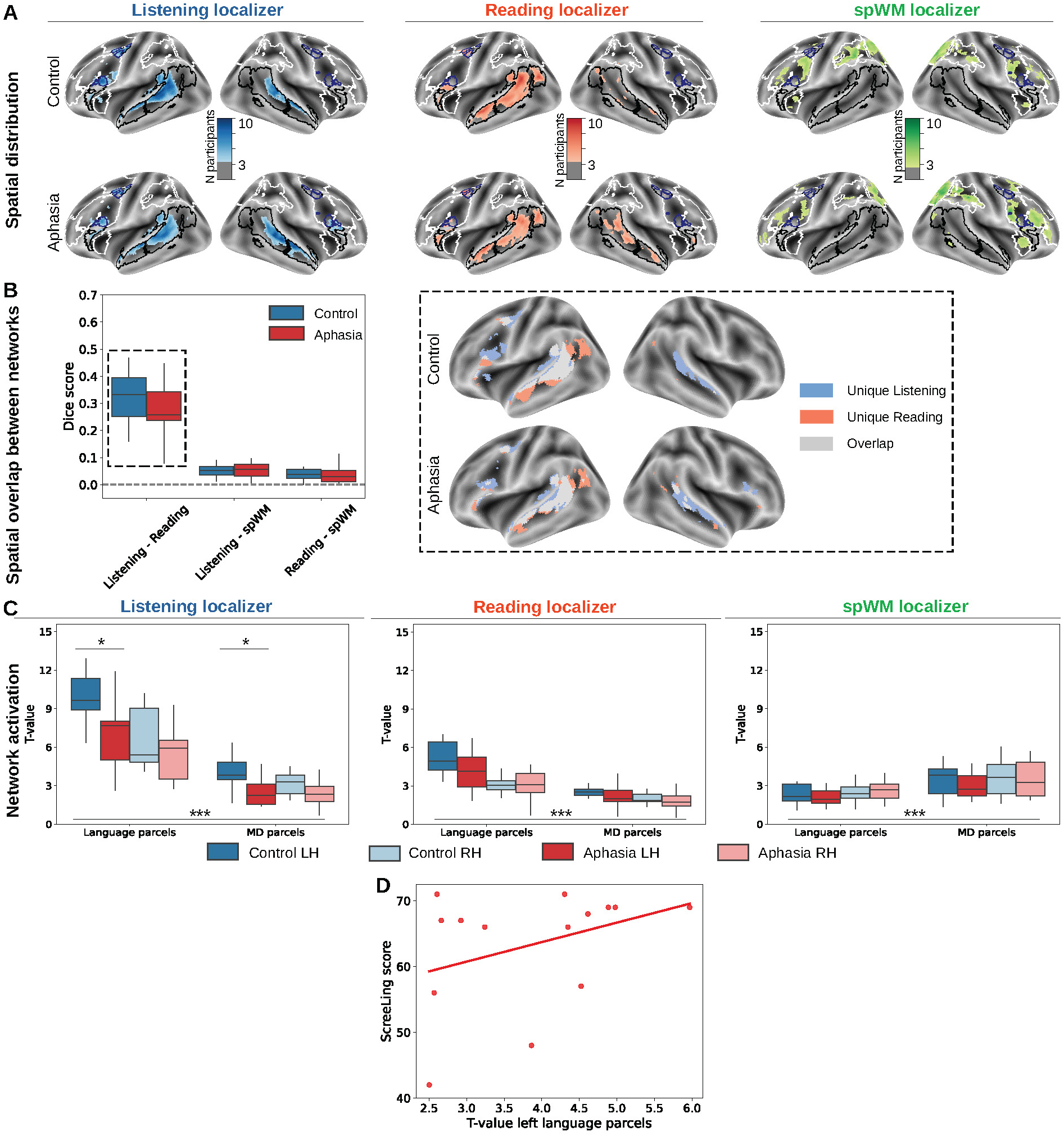
Spatial distribution and T-value activation of the localizer tasks. A. Overlap map of the single-subject-selected voxels, applying a 20% overlap threshold. Anatomical language parcels are demarcated in black, MD parcels in white, and blue demarcations indicate regions belonging to both language and MD parcels (specifically, parts of the MFG and IFG). **B.** Spatial overlap analysis using the Dice scores. Spatial overlap was observed between the listening and reading localizer task, and this overlap is visualized on the right panel (applying a 20% overlap threshold). **C.** Average T-values for controls and IWA in the predefined parcels (outlined in white in panel A) and MD (outlined in black in panel A) parcels. T-values were compared between parcels (using a linear mixed-effects model, see Methods eq. 2) and between groups (post-hoc Wilcoxon rank-sum tests). *=p<.05, ***=p<.001. **D.** Scatterplot with regression line between the T-values in the language parcels and aphasia severity (ScreeLing score) in the aphasia group. This plot shows the raw relationship between both variables (uncorrected for the confounding variables).

The listening localizer revealed overlap most present in the predefined language parcels (black masks Figure 3A). Table 1, summarizing the distribution of the selected voxels across the language and MD parcels, shows that the vast majority of selected voxels were part of the predefined language parcels for both controls and IWA. Largest overlap across participants was found in the bilateral posterior and anterior temporal lobes (see Figure 3A). For the reading localizer, the majority of selected voxels were situated within the predefined language parcels (see Table 1), with the largest overlap observed in the left posterior temporal lobe and left angular gyrus. These patterns were observed in both groups.

The spatial overlap of the listening and reading network within participants was analyzed and summarized in Figure 3B. The average Dice score was 0.32 (*±*0.11) for controls and 0.27 (*±*0.11) for IWA. No significant group difference in Dice scores was observed (Wilcoxon rank-sum test: W=54, p=0.22). The primary areas of spatial overlap between the networks were concentrated in the middle parts of the left posterior and anterior temporal lobes, largely resembling the left middle temporal gyrus. Concerning unique patterns, the listening localizer revealed a more bilateral network in the temporal lobe, with a larger overlap across participants in the superior temporal gyrus. In contrast, the reading network exhibited a more left-lateralized spatial distribution, with a larger overlap observed in the left anterior temporal lobe and left angular gyrus. These patterns hold true for both groups (see Figure 3B).

The spWM localizer task was used for MD network localization. The overlap map, shown in Figure 3A, reveals a widespread spatial distribution across frontal and parietal brain areas in both groups. The vast majority of selected voxels were situated within the predefined MD parcels for both groups (refer to Table 1).

A spatial overlap analysis revealed low Dice scores between the spWM and the listening network (0.05*±*0.03 for controls, 0.05 *±*0.03 for IWA) and between the spWM and the reading network (0.04 *±*0.02 for controls, 0.04 *±*0.03 for IWA). No significant group differences were observed for either overlap analysis (spWM - listening: W=94, p=0.91; spWM - reading: W=61, p=0.57). After applying the minimal overlap criterion of 20% of participants, no shared brain areas were detected.

#### Network activation in anatomical parcels

Next, we investigated differences in T-values between groups and between the anatomical parcels (i.e., left language, right language, left MD, right MD parcels). First, we investigated differences in T-values between parcels, irrespective of group and hemisphere (see eq. 2). T-values were higher in the language parcels for the listening localizer (F=175.04,p<.001) and for the reading localizer (F=91.68,p<.001) compared to the MD parcels. Conversely, T-values were higher in the MD parcels for the spWM localizer (F=101.47,p<.001).

Group comparisons were conducted using Wilcoxon rank-sum tests. In the listening localizer, controls exhibited higher T-values in the left language parcels (W=43, p=0.04) and the left MD parcels (W=40, p=0.03) compared to IWA, but not in the right language parcels (W=75, p=0.32) or the right MD parcels (W=60, p=0.12). An exploratory analysis in all isolated parcels (without correcting for multiple comparisons) revealed lower T-values for IWA in the left anterior temporal cortex (W=49, p=0.03), the left posterior temporal cortex (W=44, p=0.01), the left MFG (W=45, p=0.01), and the right MFG (W=48, p=0.02). These results are visualized in Supplementary Materials Figure 3.

For the reading localizer, no significant group differences were observed (left language: W=45, p=0.32; right language: W=79, p=0.94; left MD: W=48, p=0.32; right MD: W=59, p=0.42). In the isolated parcels, IWA displayed lower T-values in the left posterior temporal cortex (W=37, p=0.03) and the left angular gyrus (W=34, p=0.02) (uncorrected, see Supplementary Figure 3).

For the spWM localizer, no significant group differences were observed (left language: W=74, p=0.72; right language: W=95, p=0.87; left MD: W=68, p=0.60; right MD: W=88, p=0.87). In the isolated parcels, IWA exhibited lower T-value activation in the left insula (W=42, p=0.02), the left precentral gyrus (W=48, p=0.04), and the left anterior parietal cortex (W=37, p=0.008) (uncorrected, see Supplementary Figure 3).

#### Link to behavioral results

For the listening localizer, no significant correlation was observed between aphasia severity (ScreeLing test) and T-value activation (left language: *ρ*=0.18, p=0.50; right language: *ρ*=0.03, p=0.82; left MD: *ρ*=-0.01, p=0.93; right MD: *ρ*=-0.19, p=0.50). In the reading localizer, a significant correlation was found for the left language parcels (*ρ*=0.48, p=0.004, see scatterplot of values non-corrected for third variables in Figure 3D), but not for the other parcels (right language: *ρ*=-0.17, p=0.38; left MD: *ρ*=0.33, p=0.06; right MD: *ρ*=-0.28, p=0.12). No significant correlations were observed for the spWM localizer task (left language: *ρ*=0.04, p=0.99; right language: *ρ*=0.02, p=0.99; left MD: *ρ*=-0.10, p=0.69; right MD: *ρ*=0.13, p=0.69).

### 3.2 Analysis 2: subject-specific MD network activation during the language localizer tasks

Subsequently, we explored the involvement of subject-specific MD networks in language processing. The subject-specific MD network was defined as the top 10% most active voxels across MD parcels during the spWM task. These selected voxels served as ROIs during the listening and reading localizer tasks, where we assessed whether the T-values were significantly higher than 0 (i.e., whether the MD network is engaged during language processing). Since Analysis 1 demonstrated the primary activation of the predefined MD parcels during the spWM task (see Figure 3 and Table 1), we confined this analysis to the MD parcels. The results for this analysis are visualized in Figure 4.

**Figure 4.**
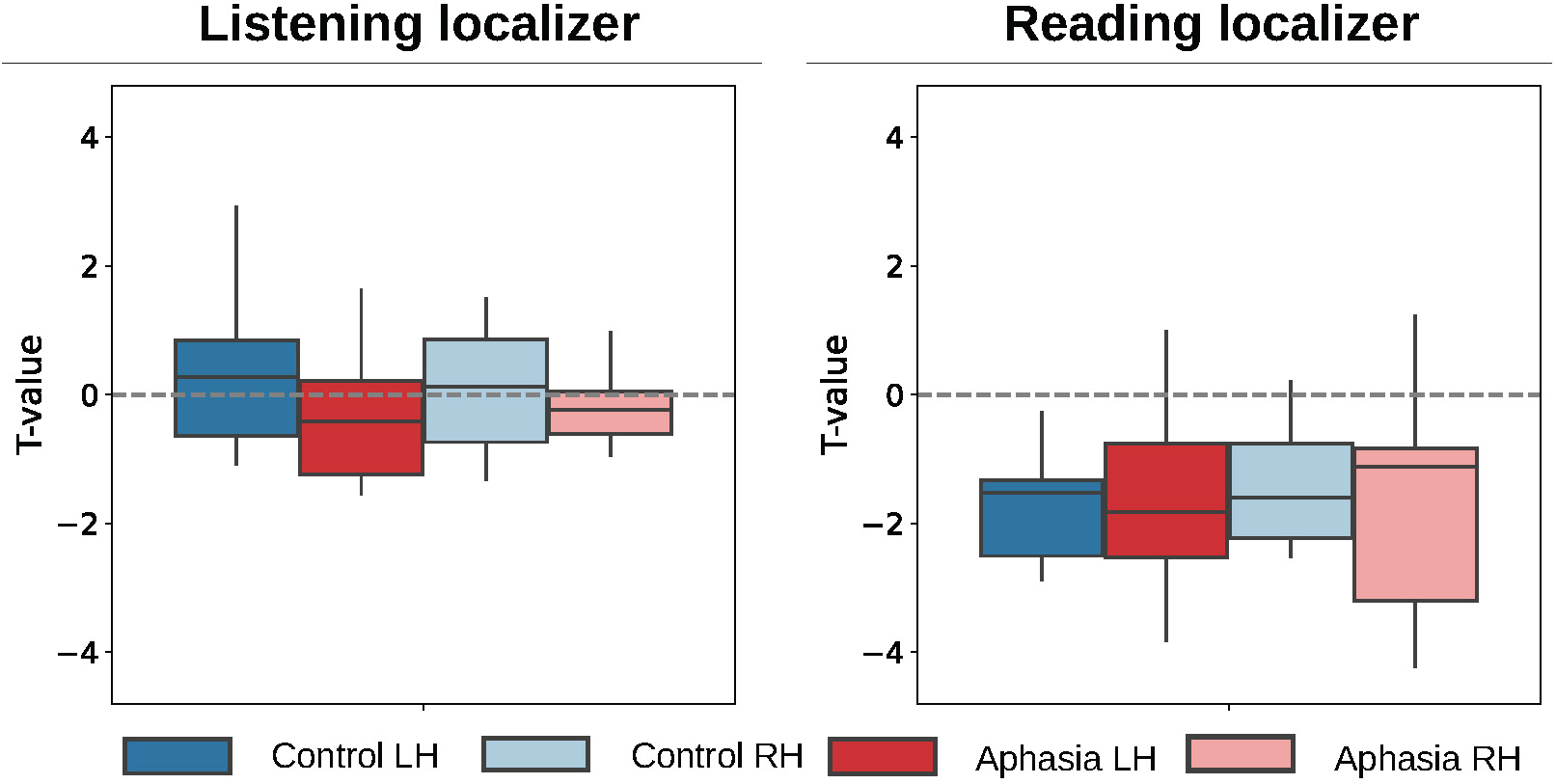
Single-subject MD activation during language processing. Average T-values of single- subject-defined MD network. Voxels were selected as the top 10% most active voxels (applying T-value and cluster thresholding) across all MD parcels (per hemisphere) during the spWM localizer task, and used as ROIs during the listening and reading localizer task. T-values were compared to 0 using one-sided Wilcoxon signed-rank tests. *=p<.05, **=p<.01, ***=p<.001.

No significant activation was observed in the listening localizer for either group or hemisphere (controls, left MD: V=58, p=0.42; controls, right MD: V=50, p=0.69; IWA, left MD: V=28, p=0.94; IWA, right MD: V=36, p=0.94). For the isolated parcels, an exploratory analysis (not controlling for multiple comparisons) revealed activation higher than 0 for controls in the left (V=84, p=0.002) and right opercular part of the IFG (V=75, p=0.02), the left (V=84, p=0.002) and right precentral gyrus (V=79, p=0.009) and the left (V=87, p<.001) and right insula (V=91, p<.001). However, IWA did not exhibit significant activation in isolated parcels. The results of this exploratory analysis in isolated parcels are visualized in Supplementary Materials, Figure 4.

For the reading localizer, no significant activation was detected (controls, left MD: V=0, p=1; controls, right MD: V=1, p=1; IWA, left MD: V=10, p=1; IWA, right MD: V=12, p=1). For both groups, we did not observe any significant activation in the isolated parcels (see Supplementary Figure 4).

Since these values do not signify meaningful activation, we refrained from conducting group comparisons or correlation analyses with aphasia severity at the behavioral level.

#### Analysis 3: Reliability of selected networks

Finally, we investigated the reliability of selected functional networks using an across-task cross-validation approach. This analysis aimed to investigate whether selected networks generalize over different task modalities. The results are visualized in Figure 5.

**Figure 5.**
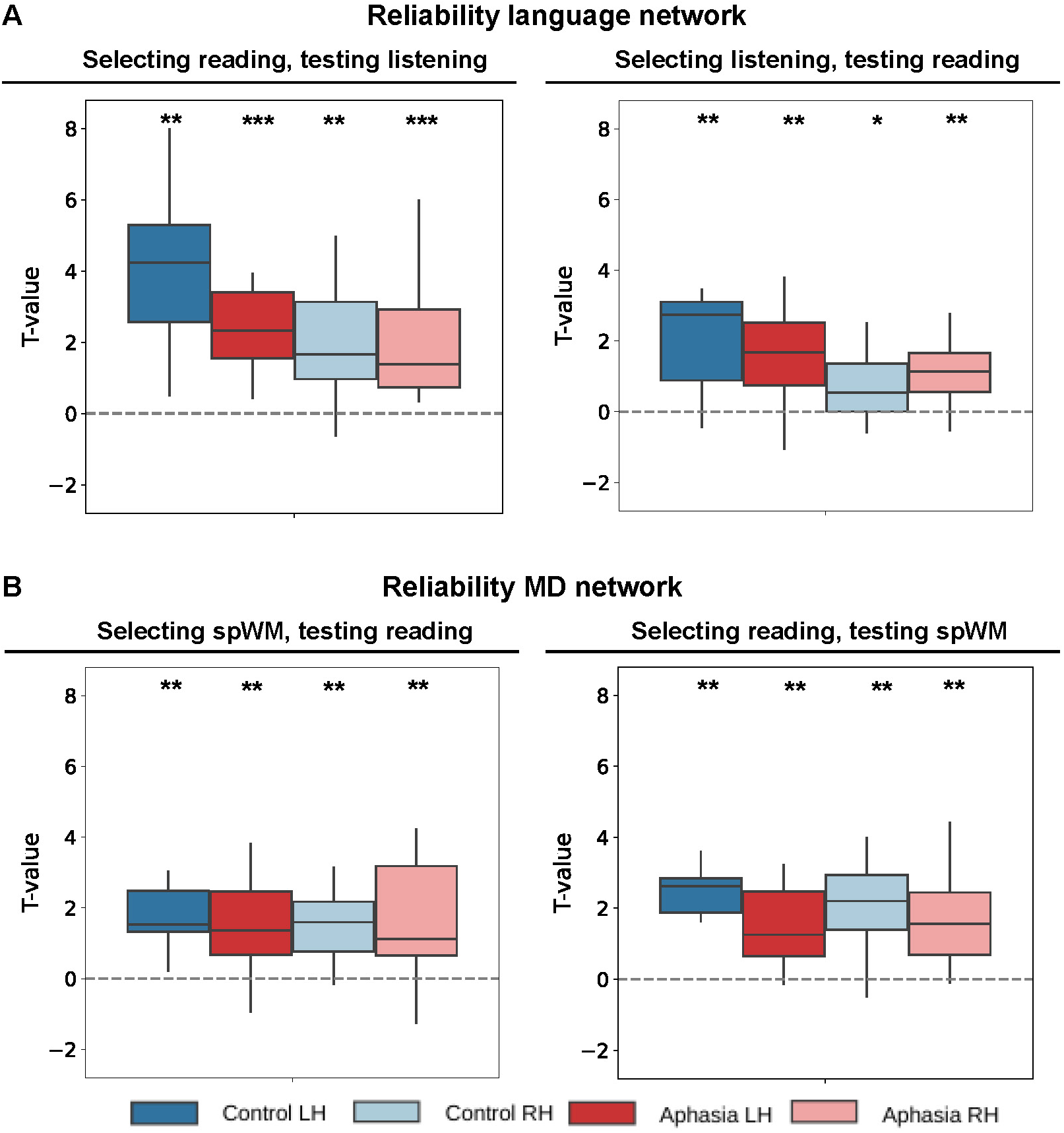
Reliability analysis using across-task cross-validation. A. Reliability of the language network, selecting based on one task and testing in another and vice-versa. **B.** Reliability of the MD network. For both panels, average T-values per participant are displayed. Voxels were selected as the top 10% most active voxels (applying T-value and cluster thresholding) across all language (panel A) and MD (panel B) parcels per hemisphere. T-values were compared to 0 using one-sided Wilcoxon signed-rank tests.

For the language network, we first selected individual voxels during the reading localizer (contrast “sentences>non-words”) using T-value and cluster thresholding across all language parcels and tested activation in these voxels for the listening localizer (contrast “intact>degraded”) against 0. These results are shown in Figure 5A. Healthy controls showed significant activation in these voxels for the left (V=66, p=0.001) and right hemisphere (V=63, p=0.003). Significant activation was also observed for IWA in the left (V=105, p<.001) and right hemisphere (V=105, p<.001). Similarly, all tests showed significant results for the reverse investigation (i.e., selecting based on the listening localizer and testing the reading localizer), visualized in Figure 5A: Controls left (V=64, p=0.002), controls right (V=56, p=0.021), IWA left (V=99, p=0.002) and IWA right (V=94, p=0.004).

Finally, we assessed the reliability and generalizability of the MD network by first selecting individual voxels during the spWM localizer task (contrast “hard>easy) using T-value and cluster thresholding across all MD parcels and testing its activation during the reading localizer (contrast “non-words>sentences”). Results are displayed in Figure 5B. Healthy controls showed significant activation in these voxels for the left (V=66, p=0.001) and right hemisphere (V=65, p=0.002). Significant activation was also observed for IWA in the left (V=81, p=0.006) and right hemisphere (V=79, p=0.009). Furthermore, all tests showed significant results for the reverse investigation (i.e., selecting based on the reading localizer and testing activation during the spWM localizer), visualized in Figure 5B: Controls left (V=66, p=0.001), controls right (V=64, p=0.002), IWA left (V=90, p=0.001) and IWA right(V=88, p=0.001).

## 4 Discussion

The present study applied individualized functional localization of the language and MD network to healthy older adults and chronic aphasia. The results reveal decreased activation in left-hemispheric language regions during the language tasks for IWA, with lower activation values in the reading task associated with poorer performance on the behavioral language test for aphasia. By contrast, we did not find evidence for MD network involvement in either of our groups, challenging prior findings in classical group-based anatomical ROI studies that have suggested a compensatory role of the MD network in aphasia. Finally, we demonstrated that functional localization of the language and MD networks generalize well across different tasks.

### Robust dissociation between the language and MD network in healthy older adults and chronic aphasia

Our results reveal a strong dissociation between the language and MD network across anatomical brain regions in chronic post-stroke and healthy older adults. In Analysis 1, both groups exhibited activation in widespread frontal and parietal brain regions during the spWM task (see Figure 3A and 3C, and Table 1), triggering MD network activation, in line with activation patterns reported in young individuals Fedorenko et al. (2013); Diachek et al. (2020). By contrast, the language localizers primarily activated traditional language regions in the temporal cortex and inferior frontal gyrus (see Figure 3A and 3C, and Table 1), largely corresponding to activation patterns observed in healthy young individuals (Fedorenko et al., 2011; Malik-Moraleda et al., 2022). The subject-specific-defined language and MD networks exhibited minimal spatial overlap, as measured by the Dice score (see Figure 3B), and no brain areas showed consistent overlap (i.e., no areas survived a 20% overlap threshold) between the two networks (see Figure 3B).

Additionally, we found no compelling evidence for subject-specific MD network activation during the language localizer tasks in Analysis 2 (see Figure 4). Nevertheless, a supplementary analysis (see Supplementary Figure 3) revealed significant activation in specific isolated MD parcels such as the bilateral insula, the bilateral precentral gyrus and the bilateral IFG for the control group only. It is crucial to note that these results 1) further speak against a compensatory role for IWA in language, as the activation was only observed in healthy controls, 2) were specific to the listening localizer and not the reading localizer, 3) emerged from an exploratory analysis without controlling for multiple comparisons and 4) involved MD region selection in isolated, small parcels without T-value or cluster thresholding, possibly reflecting selection of singular or noisy MD voxels. When applying a robust searching approach using T-value and cluster thresholding across parcels, selecting the most consistently activated voxels, no significant activation was observed for either group in either language task (see Figure 4).

In contrast to prior studies, our results argue against MD network compensation in post-stroke aphasia (Geranmayeh et al., 2014, 2017; Brownsett et al., 2014; Allendorfer et al., 2012; Griffis et al., 2017b). However, interpretation of prior results is complicated, as indicated in a recent meta-analysis by Wilson and Schneck (2021). First, many studies did not compare activation patterns in IWA to healthy controls. Consequently, it is unknown whether the observed activation reflects a true compensation mechanism in IWA or rather baseline activity required to perform the task (Wilson and Schneck, 2021). Additionally, regions within the MD network that were linked to aphasia severity, like the prefrontal cortex and supplementary motor area (Allendorfer et al., 2012; Griffis et al., 2017b), are often active in healthy individuals as well during language tasks accompanied with similar cognitive demands (e.g., semantic decision tasks) (Wilson and Schneck, 2021). Finally, prior studies used traditional group-based ROI averaging approaches, which assume equal functions for all participants in a certain anatomical ROI. However, there is substantial variability across participants in both anatomy and function of the brain (Juch et al., 2005; Fedorenko et al., 2010, 2012; Blank et al., 2017), and the language and MD networks can lie in close proximity to each other while activating distinct subregions of the same anatomical brain region (e.g., in the inferior frontal gyrus and Broca’s area) (Fedorenko et al., 2012). In line with this, our results demonstrated activation in anatomical MD parcels during our language localizer tasks as well (see activation in MD parcels for the listening and reading localizer in Figure 3C, or isolated parcels in Supplementary Figure 3), but no such language activation was found in MD regions that were subject-specific defined (i.e., subject-specific activation patterns during the spWM task).

However, offering a nuanced interpretation of our findings is essential. The present study investigated the involvement of the MD network in chronic post-stroke aphasia during core language functions, i.e., passive listening and passive reading. Whether the MD network is involved and compensates for aphasia in more complex language tasks accompanied with cognitive task demands known to activate the MD network in healthy young individuals (Diachek et al., 2020), remains a subject for further investigation. Similarly, exploring MD network activation during natural conversations, involving short-term memory storage of relevant information and speech-motor planning, raises intriguing research questions in the context of aphasia. In addition, assessing the role of the MD network in more severe cases of aphasia or during the acute stage post-stroke requires future investigation. Nevertheless, conducting functional localization approaches for these populations poses a challenge, as individuals must execute more complex cognitive tasks triggering MD network activation in the scannera task complicated by the fact that 88% of IWA in the acute stage also contend with concurrent cognitive issues (El Hachioui et al., 2014). Consequently, our results cannot be generalized to all aphasia patients and overall language functioning, but rather pertain specifically to the chronic stage and passive, receptive language functions.

### Normalized activity of the left-hemispheric language network supports behavioral language performance

Focusing on the language network, we found decreased activation for IWA compared to controls during the language localizer tasks, with most effects specific to the left hemisphere. In the listening localizer, IWA exhibited decreased activity across left-hemispheric parcels, but no group differences were found across parcels for the reading localizer. Nonetheless, results for the reading localizer show a trend towards lower values in IWA compared to healthy controls across left-hemispheric language parcels (see Figure 3C), with lower values in these parcels associated with poorer language performance on an aphasia test battery (see Figure 3D). In isolated parcels, an exploratory analysis revealed decreased activity for IWA in the left anterior and posterior temporal cortex and bilateral MFG for the listening localizer, and decreased activity in the left posterior temporal cortex and angular gyrus during the reading localizer. However, caution is warranted in interpreting results in isolated parcels as they were not corrected for multiple comparisons.

Overall, these results support general findings in the literature showing decreased left-hemispheric language activity (Li et al., 2022; Wilson and Schneck, 2021), and that normalization of this activity benefits language performance (Li et al., 2022; Wilson and Schneck, 2021). By contrast, the role of the right hemisphere in language recovery in aphasia is a subject of longstanding debate (Price and Crinion, 2005; Heiss and Thiel, 2006; Crosson et al., 2007; Li et al., 2022; Wilson and Schneck, 2021). While our study did not find evidence supporting compensatory right-hemispheric mechanisms in aphasia, prior research suggests that right-hemispheric recruitment may depend on factors such as lesion characteristics, individual recovery patterns, and the specific language components assessed during scanning (Jarso et al., 2013; Sebastian et al., 2016; Li et al., 2022; Stefaniak et al., 2022). Additionally, understanding the role of spared left-hemispheric tissue in language recovery requires considering inter-individual variability and comparing to baseline activity in healthy controls. A recent study found no support for the perilesional neuroplastic recruitment hypothesis in aphasia, i.e., no enhanced activity in brain areas surrounding individual lesions, but some evidence supporting recruitment of ipsilesional brain regions distant to the lesion (DeMarco et al., 2022). Further studies employing individualized approaches, treating individual differences as a variable of interest, are imperative in understanding language network reorganization in aphasia (Seghier and Price, 2018).

Our results further revealed no discernible group differences in MD network activity during the spWM localizer task (see 3C), which aligns with our results at the behavioral level (see Methods section for statistics). Nevertheless, group differences were found in isolated MD parcels, although these results emerged from an exploratory analysis not controlling for multiple comparisons and not applying T-value and cluster thresholding (see Analysis 1 and Supplementary Figure 3). Furthermore, we found no link between MD network activation and aphasia severity (Analysis 1). Although it is well-established that cognition and working memory are frequently impaired in aphasia and can impact language recovery (El Hachioui et al., 2014; Salis et al., 2015; Murray et al., 2018), previous studies have indicated that these comorbid cognitive issues depend on lesion characteristics and the experimental task at hand (Salis et al., 2015; Murray et al., 2018; Kasselimis et al., 2018). Since the language and MD network can lie in close proximity to each other (Fedorenko et al., 2012), it is most likely that the MD network can be damaged in IWA depending on individual lesion characteristics. Therefore, future research may delve deeper into the role of the MD network in language recovery across a diverse range of cognitive tasks and modalities, extending beyond spatial working memory as employed in our study.

### Reliability and generalizability of functional networks

Our results further demonstrate that single-subject network localization generalizes across different tasks using an across-task cross-validation approach (see Figure **??**), showing that our method can reliably detect functional networks within participants. However, it is important to note that our investigation tested activation against 0 at the group level, while certain participants did not display T-values larger than 0 (see boxplots in Figure **??**). We argue that specific limitations and potential improvements in functional localization can be identified, which must be addressed in future research to develop more robust localization methods for individual diagnostics.

Analysis 3 assessed the reliability of identified networks based on methodology established in prior work involving healthy young participants (e.g., see (Diachek et al., 2020; Shain et al., 2020)). While this analysis demonstrates cross-validation of T-values, it does not cross-validate *which* voxels to select, or in other words, which voxels actually form the single-subject functional network of interest. Commonly, researchers select the top 10% most active voxels in isolated anatomical ROI parcels. As such an approach could potentially result in the inclusion of singular, noisy voxels, we slightly adapted this procedure to make the search approach more robust. Specifically, we opted to select the top 10% most active across the union of all anatomical parcels, employing T-value and cluster thresholding. Nonetheless, we argue that future research should aim to develop more robust search approaches to define a single network per participant in a reliable manner that generalizes well over different tasks.

### Similarities and differences between the listening and reading localizer tasks

Although we have shown that the language network generalizes over task modalities, and spatial overlap was found between different localizer tasks (average Dice score of 0.32 for controls and 0.27 for IWA, see Figure 3B), differences can be observed between the obtained networks. The listening localizer exhibited larger spatial overlap across participants in bilateral temporal regions and higher overall T-values compared to the reading localizer. By contrast, the spatial overlap analysis across participants revealed stronger overlap for the reading localizer in the left middle temporal gyrus, the left anterior temporal lobe and left angular gyrus. A correlation with aphasia severity was only significant in the reading task.

Stronger left-hemispheric specificity for the reading network and more bilateral, widespread activation for the listening network align with prior work comparing both modalities in young individuals (Buchweitz et al., 2009). However, differences in contrast condition can also explain the observed differences in outcomes. While the “sentences>non-words” contrast in the reading task filters individual phonemes and syllables, the “intact>degraded” contrast in the listening task does not. Consequently, brain regions implicated in processing phonemes and speech-like sounds, notably the STG in young individuals (Yi et al., 2019), likely exhibited relatively heightened activation during the listening task. In contrast, the reading task might have induced activation more specific to surviving higher-level language functions, which might explain the stronger correlation we found with aphasia severity.

A final noteworthy difference between both language tasks is that we used the reversed “non-words>sentences” contrast during the reading localizer to assess the generalizability of the MD network in Analysis 3, while we did not utilize the reverse contrast for the listening localizer (i.e., “degraded>intact”). Although the paper that originally introduced the listening localizer (Scott et al., 2017) suggests the “degraded>intact” contrast can also be utilized for MD network localization, we refrained from this analysis a priori for 2 main reasons. First, this reversed contrast is less frequently used in the literature to select the language network. Second, our “degraded” condition was different from the control condition of Scott et al. (2017). Specifically, our study used noise shaped with the same spectrum as the intact stimuli making speech completely incomprehensible, while the listening localizer employed by Scott et al. (2017) incorporated high-frequency noise to reduce speech intelligibility (however, phonemes and syllables can still be identified when listening attentively). Hence, we hypothesized that our “degraded” contrast would not engage the MD network, which was confirmed by Analysis 2 (i.e., no negative values in Figure 4 unlike the reading localizer). Notwithstanding our difference in contrast employed in the listening localizer, our contrast was also endorsed by Scott et al. (2017) as a control to localize the language network, who anticipated no substantial differences in network localization.

## Conclusion

This study reports a novel application of individualized functional localization to investigate the role of the MD network during language processing in chronic aphasia and healthy older adults. Our results demonstrate that the MD network does not support passive, receptive language functions in both populations, arguing against a compensatory role of the MD network for aphasia. Rather, our findings agree with the general observation in the literature that normalized activity in the left-hemispheric language network supports behavioral language outcomes. Future studies may apply individualized functional localization to a wider range of language tasks, and include larger, more diverse samples of aphasia patients.

## Data and Code Availability Statements

The pseudonymized study data and code to reproduce the figures and findings of this study will be made publicly available at the Open Science Framework upon publication. Please note that the MRI data cannot be shared under any circumstance, as lesioned MRI data are person-specific and therefore cannot be considered anonymous.

## Acknowledgements

The authors would like to express their gratitude to all the participants, particularly those with aphasia and their families that supported them. A warm thanks to prof. Evelina Fedorenko for providing her advice on the experimental protocol. We would also like to extend our thanks to Dr. Klara Schevenels and Dr. Jill Kries for their assistance in the recruitment process, as well as the individuals that helped with the data collection: Katja De Meyer, Olivia Gysels, Laura Stalmans and Emilie Vanhove.

## References

1. Accou, B., Jalilpour Monesi, M., Montoya, J., Van hamme, H., and Francart, T. (2021). Modeling the relationship between acoustic stimulus and eeg with a dilated convolutional neural network. 2020 *28th European Signal Processing Conference (EUSIPCO)*, pages 1175–1179.

2. Allendorfer, J. B., Kissela, B. M., Holland, S. K., and Szaflarski, J. P. (2012). Different patterns of language activation in post-stroke aphasia are detected by overt and covert versions of the verb generation fmri task. Med Sci Monit, 18(3):CR135–137.

3. Blank, I. A., Kiran, S., and Fedorenko, E. (2017). Can neuroimaging help aphasia researchers? addressing generalizability, variability, and interpretability. Cogn Neuropsychol, 34(6):377–393.

4. Brownsett, S. L., Warren, J. E., Geranmayeh, F., Woodhead, Z., Leech, R., and Wise, R. J. (2014). Cognitive control and its impact on recovery from aphasic stroke. Brain, 137(Pt 1):242–254.

5. Buchweitz, A., Mason, R. A., Tomitch, L. M., and Just, M. A. (2009). Brain activation for reading and listening comprehension: An fmri study of modality effects and individual differences in language comprehension. Psychol Neurosci, 2(2):111–123.

6. Crinion, J. and Price, C. J. (2005). Right anterior superior temporal activation predicts auditory sentence comprehension following aphasic stroke. Brain, 128(Pt 12):2858–2871.

7. Crinion, J. T., Warburton, E. A., Lambon-Ralph, M. A., Howard, D., and Wise, R. J. S. (2006). Listening to narrative speech after aphasic stroke: The role of the left anterior temporal lobe. Cerebral Cortex, 16(8):1116–1125.

8. Crosson, B., McGregor, K., Gopinath, K. S., Conway, T. W., Benjamin, M., Chang, Y.-Y., Moore, A. B., Raymer, A. M., Briggs, R. W., Sherod, M. G., Wierenga, C. E., and White, K. D. (2007). Functional mri of language in aphasia: A review of the literature and the methodological challenges. Neuropsychol Rev, 17(2):157–177.

9. DeMarco, A. T., van der Stelt, C., Paul, S., Dvorak, E., Lacey, E., Snider, S., and Turkeltaub, P. E. (2022). Absence of perilesional neuroplastic recruitment in chronic poststroke aphasia. Neurology, 99(2):e119–e128.

10. Diachek, E., Blank, I., Siegelman, M., Affourtit, J., and Fedorenko, E. (2020). The domain-general multiple demand (md) network does not support core aspects of language comprehension: A large-scale fmri investigation. Journal of Neuroscience, 40(23):4536–4550.

11. Duncan, J. (2010). The multiple-demand (md) system of the primate brain: Mental programs for intelligent behaviour. Trends Cogn Sci, 14(4):172–179.

12. El Hachioui, H., Visch-Brink, E. G., de Lau, L. M., van de Sandt-Koenderman, M. W., Nouwens, F., Koudstaal, P. J., and Dippel, D. W. (2017). Screening tests for aphasia in patients with stroke: a systematic review. Journal of Neurology, 264(2):211–220.

13. El Hachioui, H., Visch-Brink, E. G., Lingsma, H. F., Van De Sandt-Koenderman, M. W., Dippel, D. W., Koudstaal, P. J., and Middelkoop, H. A. (2014). Nonlinguistic cognitive impairment in poststroke aphasia: A prospective study. Neurorehabilitation and Neural Repair, 28(3):273–281.

14. Fedorenko, E., Behr, M. K., and Kanwisher, N. (2011). Functional specificity for high-level linguistic processing in the human brain. Proc Natl Acad Sci U S A, 108(39):16428–33.

15. Fedorenko, E. and Blank, I. A. (2020). Broca’s area is not a natural kind. Trends Cogn Sci, 24(4):270–284.

16. Fedorenko, E., Duncan, J., and Kanwisher, N. (2012). Language-selective and domain-general regions lie side by side within broca’s area. Current Biology, 22(21):2059–2062.

17. Fedorenko, E., Duncan, J., and Kanwisher, N. (2013). Broad domain generality in focal regions of frontal and parietal cortex. Proceedings of the National Academy of Sciences, 110(41):16616–16621.

18. Fedorenko, E., Hsieh, P.-J., Nieto-Castañón, A., Whitfield-Gabrieli, S., and Kanwisher, N. (2010). New method for fmri investigations of language: Defining rois functionally in individual subjects. Journal of Neurophysiology, 104(2):1177–1194.

19. Fridriksson, J. (2010). Preservation and modulation of specific left hemisphere regions is vital for treated recovery from anomia in stroke. J Neurosci, 30(35):11558–11564.

20. Fridriksson, J., Baker, J. M., and Moser, D. (2009). Cortical mapping of naming errors in aphasia. Hum Brain Mapp, 30(8):2487–2498.

21. Friston, K., Jezzard, P., and Turner, R. (1994). Analysis of functional MRI time-series. Human Brain Mapping, 1(2):153–171.

22. Geranmayeh, F., Brownsett, S. L., and Wise, R. J. (2014). Task-induced brain activity in aphasic stroke patients: What is driving recovery? Brain, 137(Pt 10):2632–2648.

23. Geranmayeh, F., Chau, T. W., Wise, R. J. S., Leech, R., and Hampshire, A. (2017). Domain-general subregions of the medial prefrontal cortex contribute to recovery of language after stroke. Brain, 140(7):1947–1958.

24. Griffis, J. C., Nenert, R., Allendorfer, J. B., and Szaflarski, J. P. (2017a). Linking left hemispheric tissue preservation to fmri language task activation in chronic stroke patients. Cortex, 96:1–18.

25. Griffis, J. C., Nenert, R., Allendorfer, J. B., Vannest, J., Holland, S., Dietz, A., and Szaflarski, J. P. (2017b). The canonical semantic network supports residual language function in chronic post-stroke aphasia. Hum Brain Mapp, 38(3):1636–1658.

26. Hartwigsen, G. and Saur, D. (2019). Neuroimaging of stroke recovery from aphasia - insights into plasticity of the human language network. Neuroimage, 190:14–31.

27. Heiss, W.-D. and Thiel, A. (2006). A proposed regional hierarchy in recovery of post-stroke aphasia. Brain Lang, 98(1):118–123.

28. Huygelier, H., Schraepen, B., Demeyere, N., and Gillebert, C. R. (2019). The Dutch version of the Oxford Cognitive Screen (OCS-NL): normative data and their association with age and socio-economic status. *Aging*, Neuropsychology, and Cognition, 27(5):765–786.

29. Jarso, S., Li, M., Faria, A., Davis, C., Leigh, R., Sebastian, R., Tsapkini, K., Mori, S., and Hillis, A. E. (2013). Distinct mechanisms and timing of language recovery after stroke. Cogn Neuropsychol, 30(7-8):454–475.

30. Johnson, L., Basilakos, A., Yourganov, G., Cai, B., Bonilha, L., Rorden, C., and Fridriksson, J. (2019). Progression of aphasia severity in the chronic stages of stroke. American Journal of Speech-Language Pathology, 28(2):639–649.

31. Juch, H., Zimine, I., Seghier, M. L., Lazeyras, F., and Fasel, N. J. (2005). Anatomical variability of the lateral frontal lobe surface: Implication for intersubject variability in language neuroimaging. Neuroimage, 24(2):504–514.

32. Kasselimis, D., Angelopoulou, G., Simos, P., Petrides, M., Peppas, C., and Velonakis, Georgios, e. a. (2018). Working memory impairment in aphasia: The issue of stimulus modality. J Neurolinguistics, 48:104–116.

33. Keuleers, E. and Brysbaert, M. (2010). Wuggy: A multilingual pseudoword generator. Behavior Research Methods, 42(3):627–633.

34. Kries, J., De Clercq, P., Lemmens, R., Francart, T., and Vandermosten, M. (2023). Acoustic and phonemic processing are impaired in individuals with aphasia. Scientific Reports, 13(1):11208.

35. LaCroix, A. N., James, E., and Rogalsky, C. (2021). Neural resources supporting language production vs. comprehension in chronic post-stroke aphasia: A meta-analysis using activation likelihood estimates. Front Hum Neurosci, 15:680933.

36. Li, R., Mukadam, N., and Kiran, S. (2022). Functional mri evidence for reorganization of language networks after stroke. Handb Clin Neurol, 185:131–150.

37. Malik-Moraleda, S., Ayyash, D., Gallée, J., Affourtit, J., Hoffmann, M., Mineroff, Z., Jouravlev, O., and Fedorenko, E. (2022). An investigation across 45 languages and 12 language families reveals a universal language network. Nature Neuroscience, 25(8):1014–1019.

38. Murray, L., Salis, C., Martin, N., and Dralle, J. (2018). The use of standardised short-term and working memory tests in aphasia research: A systematic review. Neuropsychol Rehabil, 28(3):309–351.

39. Nenert, R., Allendorfer, J. B., Martin, A. M., Banks, C., Vannest, J., Holland, S. K., Hart, K. W., Lindsell, C. J., and Szaflarski, J. P. (2018). Longitudinal fmri study of language recovery after a left hemispheric ischemic stroke. Restor Neurol Neurosci, 36(3):359–385.

40. Oldfield, R. (1971). The assessment and analysis of handedness: the edinburgh inventory. Neuropsy- chologia, 9(1):97–113.

41. Pillay, S. B., Gross, W. L., Graves, W. W., Humphries, C., Book, D. S., and Binder, J. R. (2018). The neural basis of successful word reading in aphasia. J Cogn Neurosci, 30(4):514–525.

42. Postman-Caucheteux, W. A., Birn, R. M., Pursley, R. H., Butman, J. A., Solomon, J. M., Picchioni, D., McArdle, J., and Braun, A. R. (2010). Single-trial fmri shows contralesional activity linked to overt naming errors in chronic aphasic patients. J Cogn Neurosci, 22(6):1299–1318.

43. Price, C. J. and Crinion, J. (2005). The latest on functional imaging studies of aphasic stroke. Curr Opin Neurol, 18(4):429–434.

44. Robson, H., Zahn, R., Keidel, J. L., Binney, R. J., Sage, K., and Lambon Ralph, M. A. (2014). The anterior temporal lobes support residual comprehension in wernicke’s aphasia. Brain, 137(Pt 3):931– 943.

45. Rorden, C., Bonilha, L., Fridriksson, J., Bender, B., and Karnath, H.-O. (2012). Age-specific ct and mri templates for spatial normalization. NeuroImage, 61(4):957–965.

46. Salis, C., Kelly, H., and Code, C. (2015). Assessment and treatment of short-term and working memory impairments in stroke aphasia: A practical tutorial. Int J Lang Commun Disord, 50(6):721–736.

47. Saur, D., Lange, R., Baumgaertner, A., Schraknepper, V., Willmes, K., Rijntjes, M., and Weiller, C. (2006). Dynamics of language reorganization after stroke. Brain, 129(Pt 6):1371–1384.

48. Scott, T. L., Gallée, J., and Fedorenko, E. (2017). A new fun and robust version of an fmri localizer for the frontotemporal language system. Cognitive Neuroscience, 8(3):167–176. PMID: 27386919.

49. Sebastian, R., Long, C., Purcell, J. J., Faria, A. V., Lindquist, M., Jarso, S., Race, D., Davis, C., Posner, J., Wright, A., and Hillis, A. E. (2016). Imaging network level language recovery after left pca stroke. Restor Neurol Neurosci, 34(4):473–489.

50. Seghier, M. L. and Price, C. J. (2018). Interpreting and utilising intersubject variability in brain function. Trends in Cognitive Sciences, 22(6):517530.

51. Shain, C., Blank, I. A., van Schijndel, M., Schuler, W., and Fedorenko, E. (2020). fmri reveals language- specific predictive coding during naturalistic sentence comprehension. Neuropsychologia, 138:107307.

52. Siegel, J. S., Ramsey, L. E., Snyder, A. Z., Metcalf, N. V., Chacko, R. V., Weinberger, K., Baldassarre, A., Hacker, C. D., Shulman, G. L., and Corbetta, M. (2016). Disruptions of network connectivity predict impairment in multiple behavioral domains after stroke. Proc Natl Acad Sci U S A, 113(30):E4367– E4376.

53. Sims, J. A., Kapse, K., Glynn, P., Sandberg, C., Tripodis, Y., and Kiran, S. (2016). The relationships between the amount of spared tissue, percent signal change, and accuracy in semantic processing in aphasia. Neuropsychologia, 84:113–126.

54. Stefaniak, J. D., Geranmayeh, F., and Ralph, M. A. L. (2022). The multidimensional nature of aphasia recovery post-stroke. Brain, 145(4):1354–1367.

55. Stockert, A., Wawrzyniak, M., Klingbeil, J., Wrede, K., Kümmerer, D., Hartwigsen, G., Kaller, C. P., Weiller, C., and Saur, D. (2020). Dynamics of language reorganization after left temporo-parietal and frontal stroke. Brain, 143(3):844–861.

56. Szaflarski, J. P., Allendorfer, J. B., Banks, C., Vannest, J., and Holland, S. K. (2013). Recovered vs. not-recovered from post-stroke aphasia: The contributions from the dominant and non-dominant hemispheres. Restor Neurol Neurosci, 31(4):347–360.

57. Szaflarski, J. P., Eaton, K., Ball, A. L., Banks, C., Vannest, J., Allendorfer, J. B., Page, S., and Holland, S. K. (2011). Poststroke aphasia recovery assessed with functional magnetic resonance imaging and a picture identification task. J Stroke Cerebrovasc Dis, 20(4):336–345.

58. Turkeltaub, P. E., Messing, S., Norise, C., and Hamilton, R. H. (2011). Are networks for residual language function and recovery consistent across aphasic patients? Neurology, 76(20):1726–1734.

59. Van Ewijk, E., Dijkhuis, L., Hofs-Van Kats, M., Hendrickx-Jessurun, M., Wijngaarden, M., and De Hilster, C. (2020). Nederlandse Benoem Test. Bohn stafleu van loghum.

60. van Oers, C. A., Vink, M., van Zandvoort, M. J., van der Worp, H. B., de Haan, E. H. F., Kappelle, L. J., Ramsey, N. F., and Dijkhuizen, R. M. (2010). Contribution of the left and right inferior frontal gyrus in recovery from aphasia: A functional mri study in stroke patients with preserved hemodynamic responsiveness. Neuroimage, 49(1):885–893.

61. Visch-Brink, E., Van de Sandt-Koenderman, M., and El Hachioui, H. (2010). ScreeLing. Houten: Bohn Stafleu Van Loghum.

62. Warren, J. E., Crinion, J. T., Lambon Ralph, M. A., and Wise, R. J. (2009). Anterior temporal lobe connectivity correlates with functional outcome after aphasic stroke. Brain, 132(Pt 12):3428–3442.

63. Wilson, S. M. and Schneck, S. M. (2021). Neuroplasticity in post-stroke aphasia: A systematic review and meta-analysis of functional imaging studies of reorganization of language processing. Neurobiol Lang (Camb), 2(1):22–82.

64. Yeo, B. T. T., Krienen, F. M., Sepulcre, J., Sabuncu, M. R., Lashkari, D., Hollinshead, M., Roffman, J. L., Smoller, J. W., Zöllei, L., Polimeni, J. R., Fischl, B., Liu, H., and Buckner, R. L. (2011). The or- ganization of the human cerebral cortex estimated by intrinsic functional connectivity. J Neurophysiol, 106(3):1125–1165.

65. Yi, H.-G., Leonard, M. K., and Chang, E. F. (2019). The encoding of speech sounds in the superior temporal gyrus. Neuron, 102(6):1096–1110.

