## Supplementaries for "Individualized functional localization of the language and multiple demand network in chronic post-stroke aphasia"

### Supplementary material

#### 1 Demographic information on the aphasia sample

**Supplementary Table 1.** Demographics and lesion information of the aphasia group

| ID(n=15) | age | sex | Handed-ness | Time since stroke (months) | Stroke type | Blood vessel | Lesioned hemi-sphere | Lesion size (ml) | SLT | NBT score (max=276) | Screeling score (max=72) |
|---|---|---|---|---|---|---|---|---|---|---|---|
| sub-007 | 72 | m | right | 47 | ischemia | MCA | left | 80.79 | yes | 261 | 68 |
| sub-008 | 43 | m | right | 39 | ischemia | MCA | left | 6.91 | yes | 274 | 71 |
| sub-010 | 62 | f | right | 45 | ischemia | MCA | left | 6.55 | yes | 263 | 71 |
| sub-011 | 70 | m | left | 45 | ischemia | MCA | bilateral | 46.99 | yes | 261 | 69 |
| sub-013 | 80 | m | right | 28 | ischemia | PCA | bilateral | 10.84 | yes | 269 | 66 |
| sub-014 | 70 | m | right | 22 | ischemia | MCA | left | 98.72 | yes | 265 | 67 |
| sub-015 | 73 | f | right | 47 | ischemia | MCA | left | 0.81 | no | 271 | 68 |
| sub-017 | 41 | f | left | 13 | ischemia | MCA | left | 78.25 | yes | 267 | 69 |
| sub-018 | 69 | m | right | 18 | ischemia | MCA | left | 3.95 | no | 270 | 67 |
| sub-020 | 51 | m | right | 31 | hemorrhage | MCA | left | 45.63 | yes | 217 | 57 |
| sub-022 | 74 | m | right | 121 | hemorrhage | MCA | left | 58.59 | yes | 266 | 69 |
| sub-024 | 80 | f | right | 26 | hemorrhage | MCA | left | 88.60 | yes | 236 | 48 |
| sub-025 | 74 | m | right | 17 | hemorrhage | MCA | left | 30.81 | yes | 271 | 66 |
| sub-027 | 70 | m | right | 21 | ischemia | PCA | left | 76.67 | yes | 150 | 56 |
| sub-031 | 41 | f | right | 45 | ischemia | MCA | left | 166.32 | yes | 164 | 63 |
| Total | 65 $\pm$ 14 | 10m 5f | 13 left 2 right | 38 $\pm$ 26 | 11 ischemia 4 hemorrhage | 13 MCA 2 PCA | 13 left 2 bilateral | 53.36 $\pm$ 46.27 | 13 yes 2 no | 247.00 $\pm$ 39.59 | 65 $\pm$ 6.47 |

SLT=speech-language therapy; NBT=Dutch naming test (Nederlandse Benoem Test); MCA=middle cerebral artery; PCA=posterior cerebral artery

#### 2 Localizer tasks

A visualization of the reading language localizer task is provided in Figure 1A, and in Figure 1B for the spatial working memory (spWM) localizer task. For the latter, a button response (left or right) was required as response. The language localizer tasks (both reading and listening) involved passive processing.

Figure 1C displays the accuracy and reaction time of the responses for the first 6 participants (4 controls and 2 IWA) in the spWM task (averages across trials). Technical issues in the interface between button presses and the stimulus computer resulted in the loss of behavioral response data for the rest of the sample, unfortunately. Despite this, a repeated measures ANOVA analysis (analyzing all individual trials) revealed a significant main effect for condition: participants responded less accurately (T=-2.04, p=0.04) and slower (T=7.07, p<.001) in the hard condition of the spWM task. Figure 1C illustrates the slower reaction time effect for each individual subject, strongly suggesting that participants experienced the hard condition as more cognitively demanding. The linear mixed effects model was created with the folowing formula:

$$\text{accuracy} \sim \text{condition} + (1|\text{subject}) \tag{1}$$

$$\text{reaction time} \sim \text{condition} + (1|\text{subject}) \tag{2}$$

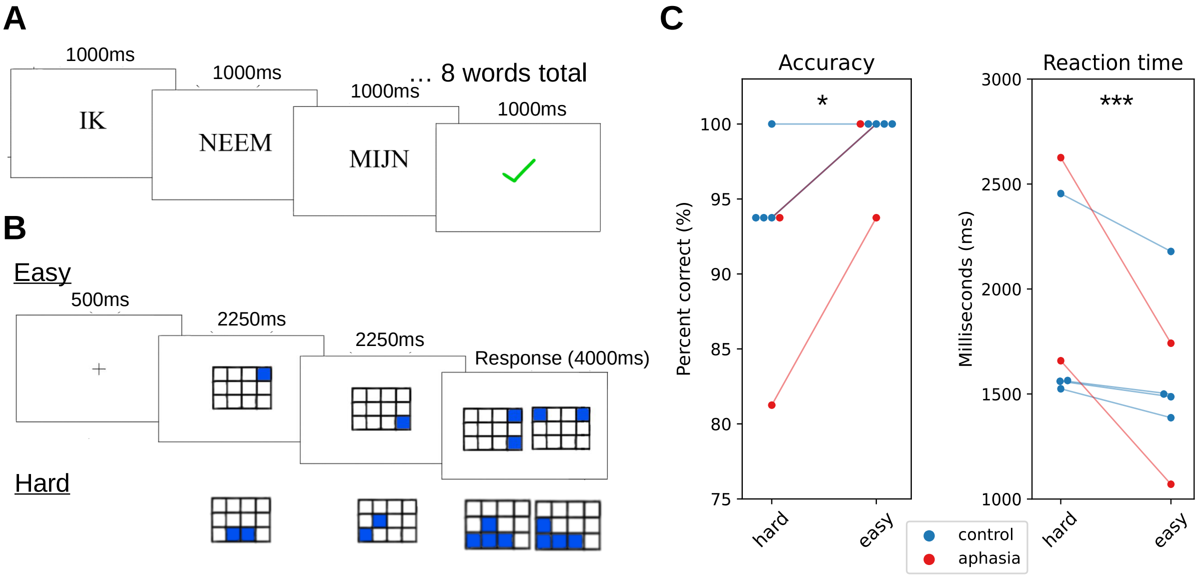


**Supplementary Fig. 1. Localizer tasks.** Illustration of the reading task (**A**) and the spatial working memory task (**B**). **C.** Average accuracies and reaction times across trials for the first 6 participants' responses in the spatial working memory task.

### 3 Predefined parcels

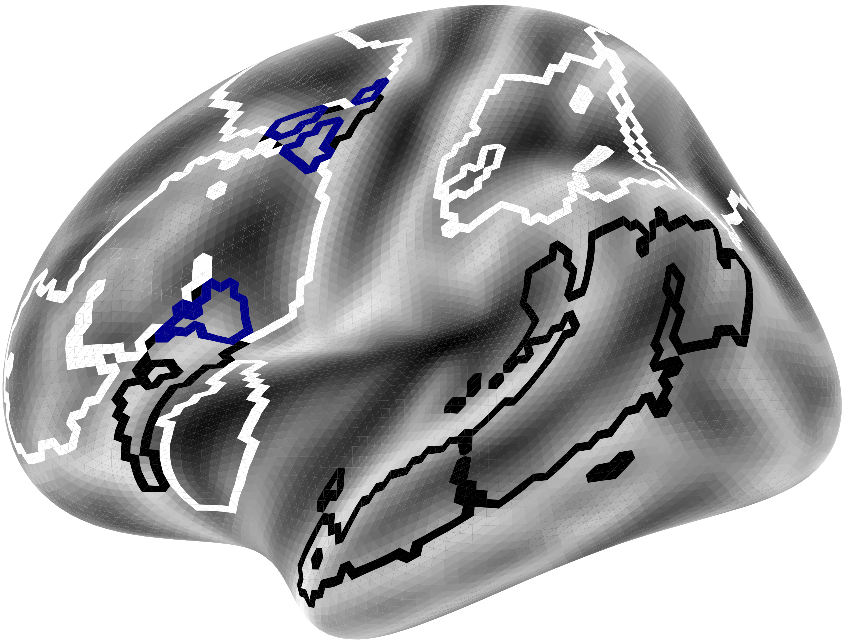

**Supplementary Fig. 2. Predefined parcels.** Black outlines indicate the language parcels, white outlines indicate the MD parcels, and blue outlines indicate regions belonging both to the language and MD parcels (present in IFG, MFG).

### 4 Network activation in individual parcels

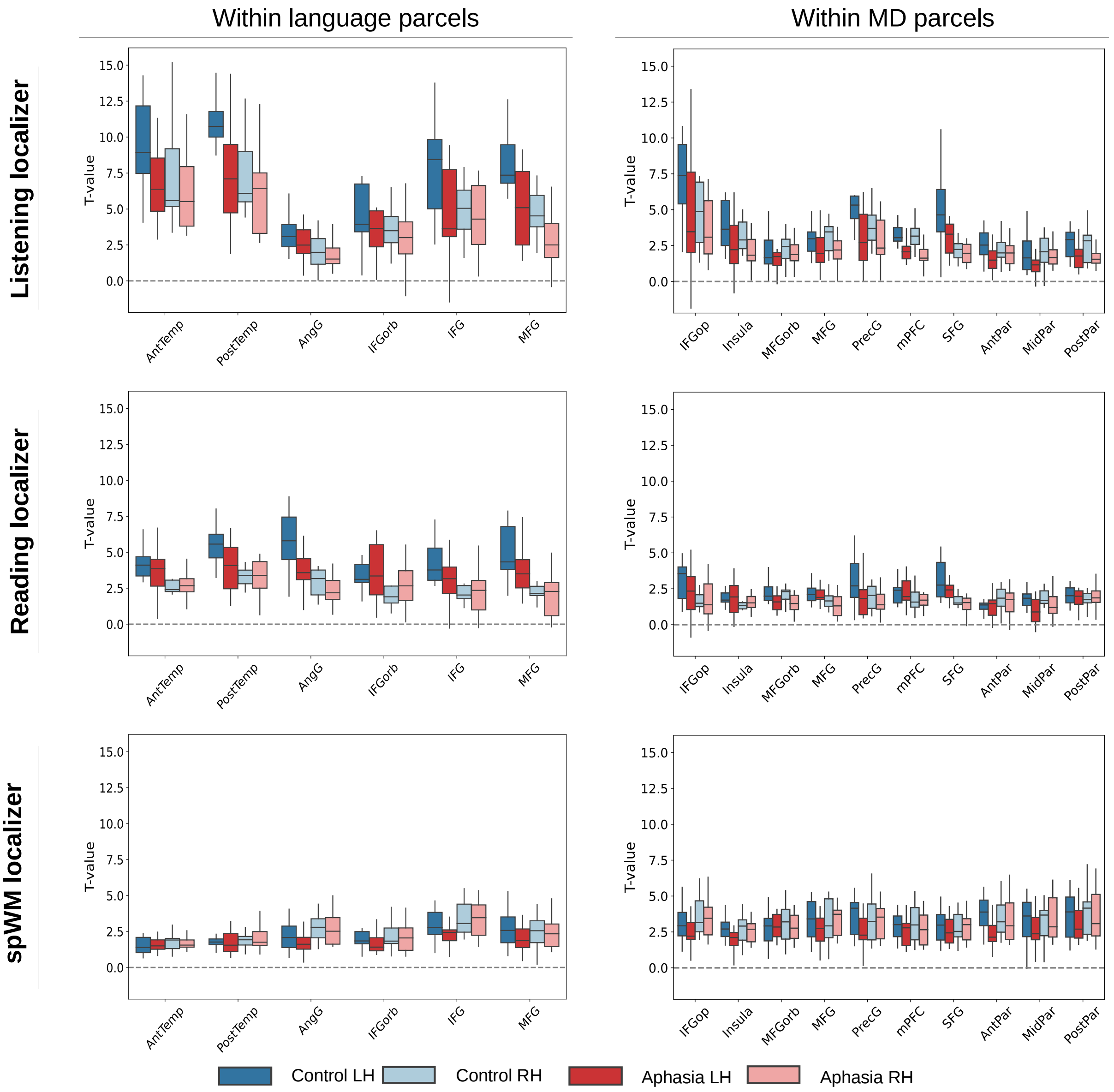


**Supplementary Fig. 3. T-values in the individual parcels for the different localizer tasks.** Voxels were selected as the top 10% most active voxels within each individual parcel. Post-hoc Wilcoxon rank-sum tests, reported in the main text, were restricted to the language parcels for the listening and reading tasks, and to the MD parcels for the spWM task. This was done since Analysis 1 indicate that these tasks primarily activated the corresponding parcels.

### 5 MD network activation in individual parcels during the language localizer tasks

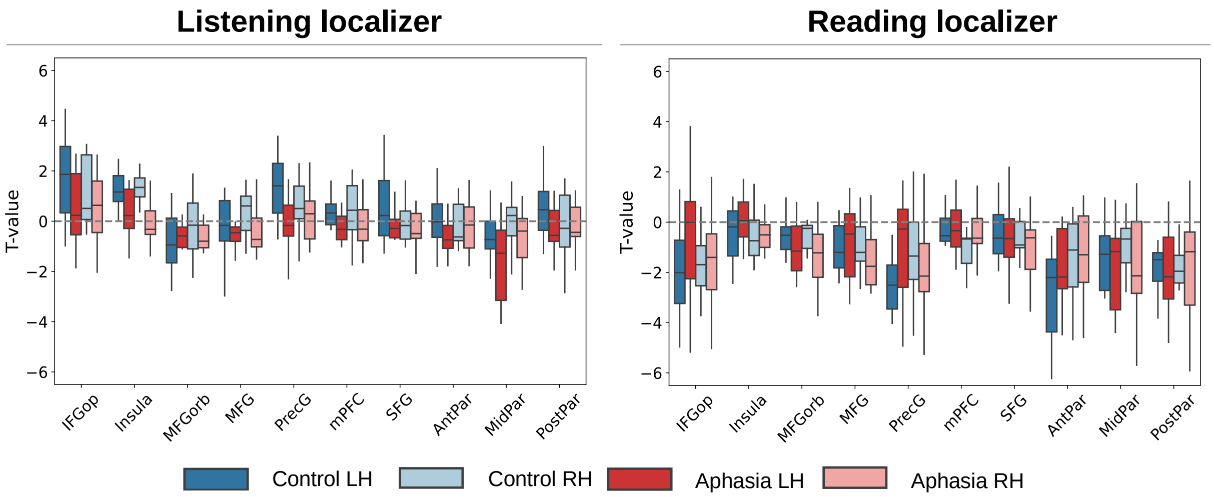


**Supplementary Fig. 4. MD network activation in the individual parcels for the listening and reading localizer task.** The MD network was single-subject-defined as the top 10% most active voxels during the spWM localizer task. These voxels were used as regions of interest during the listening and reading task. Statistics, testing whether the individual parcels exhibited T-values significantly larger than 0, are reported in the main text.
